# High frequency broadband activity detected noninvasively in infants distinguishes wake from sleep states

**DOI:** 10.1101/2025.08.08.668962

**Authors:** Ania M. Holubecki, Josset B. Yarbrough, Vinitha Rangarajan, Rachel Kuperman, Robert T. Knight, Elizabeth L. Johnson

## Abstract

High frequency broadband activity (HFB; 70–150 Hz) indexes local brain activity. It is predominantly studied using invasive measures due to signal drop off from skull attenuation. We hypothesized that HFB is detectable in infants noninvasively through fontanelles and thin skull that have not fully developed. We analyzed scalp electroencephalography (EEG) data during wake and sleep states in 19 channels from 18 infants (1–4 months, both sexes). At the group level, linear mixed-effects models revealed greater HFB power in wake versus sleep states in midline and central channels near fontanelles, as well as in occipital channels over thin skull. These differences were detected with 90% reliability using as few as 25 seconds of data per state in as few as 10 subjects. On the individual level, linear mixed-effects models revealed the same wake > sleep effect with a mean reliability of 60% when using at least 50 seconds of data per state. These findings establish that noninvasive HFB detection in infants is not only possible at sites where the skull has not fully developed, but sufficiently robust to enable systematic investigation of early cognitive development.

## Introduction

High frequency broadband activity (HFB; 70–150 Hz) indexes local population neuronal firing^1–7^. It is predominantly studied using intracranial electroencephalography (iEEG) in patients with intractable epilepsy undergoing seizure monitoring as part of their clinical care. The clinical team temporarily implants electrodes to determine the seizure onset zone and identify eloquent cortex, with the goal of resecting brain tissue to reduce epileptiform activity. iEEG studies report increases in HFB power during tasks and in wake compared to sleep states^8–10^. HFB has been difficult to detect in scalp EEG due to high-frequency signal drop off from skull attenuation and low spatial sampling^11–13^. Although scalp EEG studies have reported noninvasive HFB detection during motor tasks^14–16^, it is difficult to separate this signal from muscle noise.

iEEG in children and adolescents has enabled the study of HFB across development^17–20^. However, less is known about HFB in infancy since intracranial electrodes are rarely implanted in patients under two years of age due to thin skull hampering electrode attachment^21^. Additionally, infants have fontanelles between skull plates that have not yet fully fused^22–24^. The youngest reported patient implanted with intracranial depth electrodes was 17 months old, with a thin skull and an open anterior fontanelle^25^. The invasive monitoring was well tolerated and resulted in a successful resection of the epileptogenic zone, supporting the safety and efficacy of such procedures in very young patients. However, the case report did not include HFB. Additionally, iEEG remains a risky procedure and will likely remain rare in infants compared to older children, adolescents, and adults.

We hypothesized that the thin skull and fontanelles that increase the risk of iEEG in infants may be the environment needed to detect HFB noninvasively using scalp EEG. A study in patients undergoing decompressive hemicraniectomy, with half of their skull removed for an extended period, found that EEG signals from channels over the hemicraniectomy were more independent from each other and had enhanced task-related power at higher frequencies, including high gamma, than EEG signals from channels over the intact hemisphere^26^. In infants, channels over thin skull and fontanelles are closer to brain tissue, allowing for less signal drop off (**Figure 1**). As past intracranial work has shown that HFB is higher in wake versus sleep states^8^, we predicted that channels located over fontanelles (midline, central) and thin skull have greater HFB power in wake compared to sleep states.

**Figure 1:**
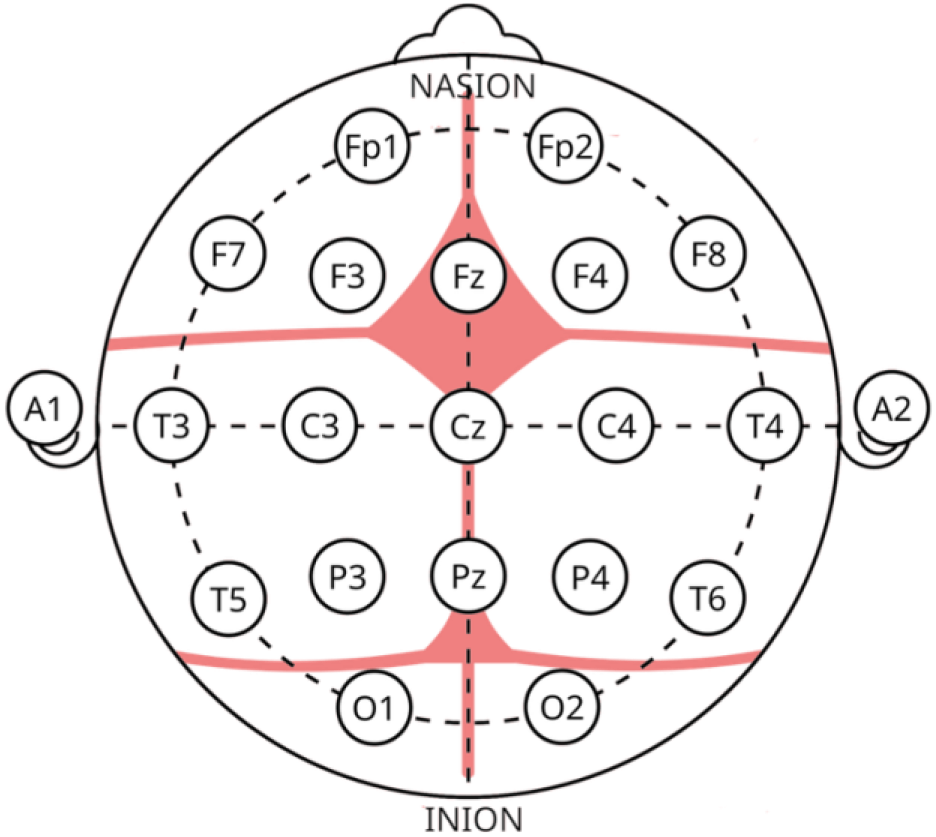
Approximate locations of the infant anterior and posterior fontanelles, as well as the frontal, coronal, sagittal, and lambdoidal sutures, in relation to scalp EEG channels arranged according to the international 10-20 system. Fontanelles and sutures are marked in pink.

## Results

### Group-level HFB differentiates between wake and sleep states with excellent reliability

At the 18-subject and all-trial group level, mixed-effects models revealed significantly greater HFB power (FDR-adjusted *p* < 0.004) during wake versus sleep states in 13 channels. The greatest differences were in midline channels Fz (*t(3211)* = 13.777), Cz (*t(3319)* = 13.265), and Pz (*t(3106) = 14*.*643*); central channels C3 (*t(2872)* = 13.410) and C4 (*t(2872)* = 12.467); and occipital channels O1 (*t(961) =* 11.370) and O2 (*t(1041) =* 17.370). Other channels with significantly greater HFB power during wake versus sleep states were Fp1 (*t(696)* = 3.545), Fp2 (*t(789)* = 11.228), P4 (*t(1748)* = 4.883), F8 (*t(2798)* = 15.135), T3 (*t(1199)* = 8.440), and T4 (*t(1305)* = 14.223). The mixed effects models also revealed significantly greater HFB power during sleep versus wake states in five channels: F3 (*t(2728)* = -13.144), F4 (*t(2624)* = -10.778), F7 (*t(1945)* = -9.768), T5 (*t(2530)* = -3.298), and T6 (*t(2591)* = -8.357). The sleep > wake difference was not significant in channel P3 (*t(2587)* = -2.878, *p* = 0.004).

When the models were run with 1–20 trials per state and replicated 10 times per trial count, reliability stabilized at around 5 trials per state, at which the majority of significant wake > sleep effects replicated at least nine times (≥90% reliability; **Figure 2**). When the models were then run with both 1–18 subjects and 1–20 trials per state, replicated 10 times per subject and trial count, reliability stabilized at around 10 subjects. Effects were detected with ≥90% reliability using as few as 10 subjects, and fewer subjects for channels O1 and O2 (**Figure 3**). When reliability was examined using an orthogonal approach as calculated by the Reliability, Effect size, And Data quality In EEG (READIE) Toolbox^27^, the internal consistency reliability for a wake > sleep HFB difference crossed the excellent threshold for midline channels Fz (M ± SD: 0.990 ± 0.008), Cz (0.979 ± 0.013), and Pz (0.993 ± 0.005); central channels C3 (0.934 ± 0.046) and C4 (0.939 ± 0.045); occipital channel O1 (0.944 ± 0.051); and channels Fp2 (0.908 ± 0.067), F8 (0.989 ± 0.007), T3 (0.998 ± 0.002), and T4 (0.969 ± 0.026). Occipital channel O2 was close to the excellent threshold (0.893 ± 0.085).

**Figure 2:**
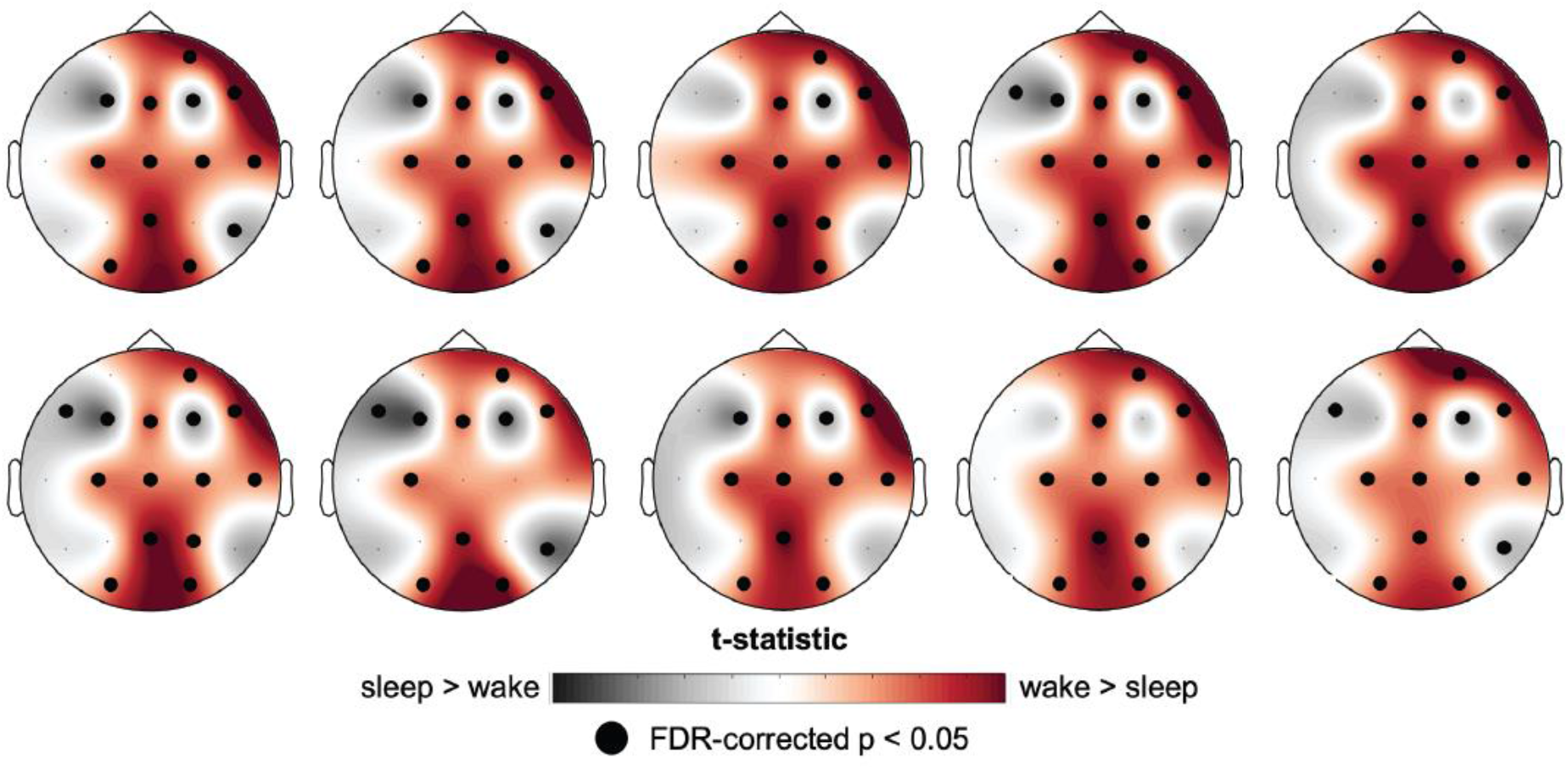
Comparison of HFB power between 5 wake trials and 5 sleep trials per subject, across 18 subjects. Each topographic plot is a replication of the per-channel mixed-effects model with the random effects of subjects and trials nested in subjects. Darker red regions of the topographic maps represent a positive t-statistic, with greater HFB power in wake versus sleep states (wake > sleep). Darker grayscale colors represent a negative t-statistic, with greater HFB power in sleep versus wake states (sleep > wake). Large black circles represent significant channels, FDR-corrected for multiple comparisons. Greater HFB power in wake versus sleep was found in midline (Fz, Cz, Pz), bilateral central (C3, C4), bilateral occipital (O1, O2) channels, and unilateral channels Fp2, F8, and T4. These results were ≥90% reliable across 10 replications.

**Figure 3:**
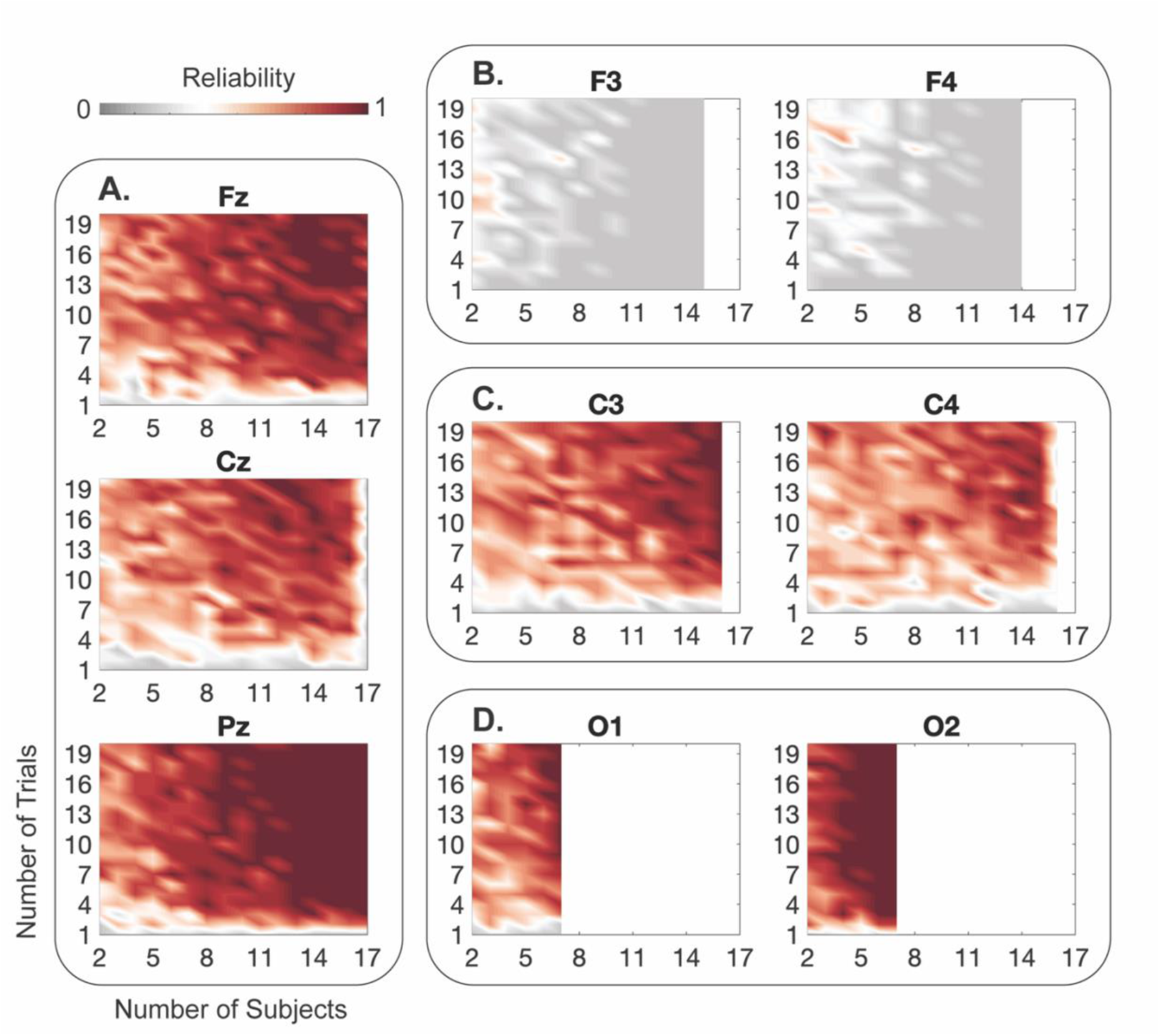
Reliability of group-level HFB power in detecting effects in representative channels, using mixed-effects models with the random effects of subjects and trials nested in subjects. Darker red colors represent greater reliability (>50%), and darker grayscale colors represent lower reliability (<50%). White on the right portions of plots in B–D represents missing data resulting from channel exclusion within individual subjects due to low data quality. The wake > sleep effect was detected with high reliability in A) midline (Fz, Cz, Pz) and C) central (C3, C4) channels when the model was run with as few as 10 subjects and 5 trials per subject, with fewer subjects required for D) occipital (O1, O2) channels. In other channels, such as B) frontal (F3, F4) channels, this effect was not reliable.

### Individual-level HFB differentiates between wake and sleep states with acceptable reliability

Having revealed that group-level HFB power is significantly greater during wake versus sleep states in channels Fz, Cz, Pz, C3, C4, O1, and O2, we calculated the reliability of detecting a wake > sleep effect in the mean HFB value across these channels at the individual level. In individual infants, the mean wake > sleep effect was detected with around 60% reliability when using 10 or more trials per state (**Figure 4**).

**Figure 4:**
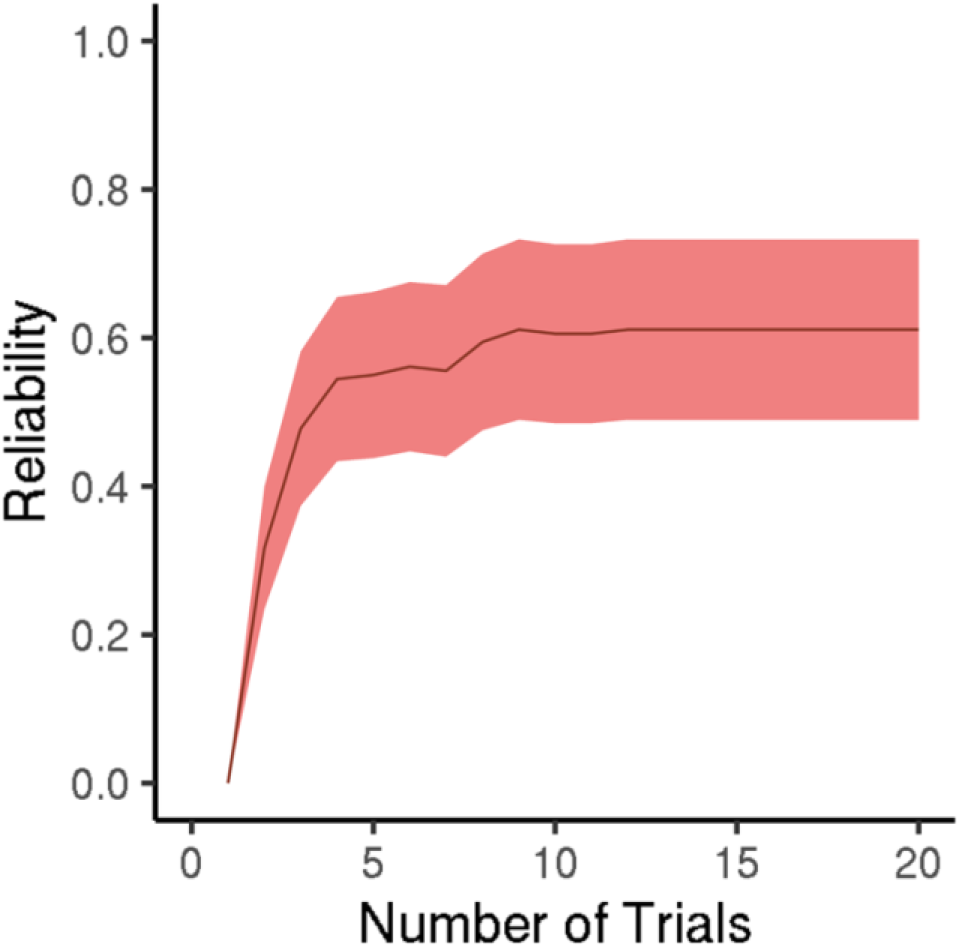
Reliability of individual-level HFB power in detecting effects using the channel mean based on group-level results (Fz, Cz, Pz, C3, C4, O1, O2) and parameters (i.e., mixed-effects model with the random effect of trial). Shading represents the standard error of the mean.

## Discussion

This is the first study to detect HFB in infants using scalp EEG. The significant wake > sleep effect characteristic of intracranial HFB is not brain region specific^8^, so the presence of these effects in some but not all channels may be influenced by characteristics of the skull. The significant wake > sleep effect in midline channels Fz, Cz, and Pz, and in central channels C3 and C4, may be detectable due to their location near the anterior fontanelle^23,28^. The significant wake > sleep effect in channels O1 and O2 may be detectable due to their location over occipital skull that is thinner than the rest of the skull^29^. Signal recorded at these channels may encounter less attenuation as it passes through less developed regions of the infant skull. Significant wake > sleep effects were reliably detected, as confirmed by two reliability measures (mostly crossing an excellent threshold of 90%, with one channel at 89% in the internal consistency reliability analysis).

These were detected using as little as 25 seconds of data in each state for each infant, in as few as 10 infants, with even fewer infants needed for occipital channels. These results establish that HFB can be studied noninvasively in infants aged 1–4 months using scalp EEG.

Although both channels Fp1 and Fp2 had a significant wake > sleep effect when using all trials and subjects, Fp1 required more data (20 trials, or 100 seconds) per state compared to other channels to reach 90% reliability, which is still a relatively small amount of data required to detect an effect. Additionally, channel F8 had a significant wake > sleep difference in HFB power when using all trials and subjects, but this effect was neither bilateral (absent in F7) nor reliably detected across 10 replications using 5 trials per state (30%). HFB differences in wake versus sleep states are not hemisphere-specific when measured intracranially^8^, so these lateralized effects are not biologically significant. Nonetheless, with an increased number of trials, these frontal electrodes may also pick up weaker signal from nearby fontanelle areas.

This study paves the way for future work to examine task-related changes in HFB activity in infancy. Infants can participate in several tasks that tap into developing cognitive processes. For example, neonates can detect grayscale and the color red, and they can detect more colors by around 6 months^30^. Additionally, the mismatch negativity (MMN) used to study prediction is evoked in infants when auditory oddball tasks are presented during sleep states^31,32^, and this paradigm elicits intracranial HFB in adults^9,33^, so it is possible that HFB differences can also be detected for oddball stimuli in infancy.

HFB power differences were also reliably detected at the individual level (60%), although not to the same extent as in group-level analyses. Reliability thresholds vary across the field and are not yet well established for developmental populations^27,34,35^. A reliability of 60% is higher than that seen in many developmental scalp EEG studies. For example, one study implemented two internal consistency reliability measures of different MMN components in toddlers aged 24–32 months^36^. They found that the early MMN had an individual level dependability ranging from .011–.650 and an individual level intraclass correlation (ICC) ranging from .002–.010. The late MMN had an individual level dependability ranging from .012–.512 and individual level ICC ranging from .001–.006. Additionally, the READIE Toolbox study reported an internal consistency reliability of around .40 on the N1 mean amplitude from their visual evoked potential task^27^. Interestingly, the same study reported that gamma activity (30-50 Hz) passed the excellent threshold of internal consistency reliability with as little as 20 seconds of data in a 2–5-month age group, whereas the 6–12-month and 13–18-month age groups passed the good threshold with the same amount of data. This difference in reliability across age groups was not present in other frequency bands, which suggests that this difference may also be due to differences in skull development. Scalp-derived HFB should be utilized with caution in a clinical setting with individual infants, but the reliability of detecting this signal with scalp EEG is better than other individual scalp EEG measures.

One limitation of our study is that we did not have exact dates of birth for each subject. This is important since fontanelles close at different stages of development. The posterior fontanelle closes around 6 to 8 weeks after birth^22^, followed by the sphenoid fontanelles around 6 months^28^, mastoid fontanelles around 6 to 18 months, and anterior fontanelle around 13 to 24 months^24^. Prior work did not find a difference in signal conductance between fontanelle and skull regions in newborns^37^, which may be due to thinner skull and tissue layers at term age compared to our 1-4 month old cohort. Another limitation is that the dataset did not include the sex and race of each subject. The anterior and posterior fontanelles are larger in Black compared to white infants^38^, and close sooner in male compared to female infants^23^. Given the spread of reliability between infants in our individual reliability analyses, it is possible that the 60% reliability at which we detected individual HFB differences could be improved by taking into consideration the age, sex, and race of the infants. Demographic information in future studies could help better define developmental periods in which HFB can be detected using scalp EEG.

64-channel caps for infants are reported to measure focal cortical activity within a few centimeters of each channel^37^. HFB is a local signal, so future studies should explore data collected from higher density caps for better specificity regarding where HFB can be detected. HFB detection in infancy provides a novel method for studying how cognitive processes emerge in early development.

## Methods

### Subjects

The study included 22 infants (age range 1–4 months, both sexes; exact ages and sex ratio unknown) undergoing outpatient scalp EEG monitoring for potential seizures at the University of California, San Francisco Benioff Children’s Hospital. All infants were determined to be seizure free during monitoring. Written informed consent was obtained from the guardians of the infants in accordance with procedures approved by the University of California, San Francisco Institutional Review Board and in accordance with the Declaration of Helsinki.

### EEG data acquisition and preprocessing

Scalp EEG data was collected using a 19-channel cap arranged according to the international 10-20 system over midline (Fz, Fpz, Cz, Pz, Oz), frontal (F7, F3, F4, F8), parietal (P3, P4), frontoparietal (Fp1, Fp2), temporal (T3, T4), central (C3, C4), and occipital (O1, O2) areas^39,40^. The data were recorded at a sampling rate of 1000 Hz. A pediatric epileptologist manually annotated the raw EEG data for sleep and wake states.

Each manually annotated data segment was split into 5-second trials, with 50% overlap between trials to maximize the amount of data retained. This resulted in a range of 42–288 sleep trials and 20–261 wake trials per subject after preprocessing. The sleep data from one subject and wake data from three subjects were excluded due to an insufficient number of trials within a state (fewer than 20). Eighteen subjects had both sleep and wake data and were analyzed.

Sleep and wake data were preprocessed separately for each subject. The raw data were bandpass (0.1–200 Hz Butterworth finite impulse response) and notch (60, 120, 180 Hz) filtered to remove line noise. Channels and trials were then automatically excluded if they exceeded two standard deviations of the mean across channels and trials. Independent component analysis (ICA) was performed; each component and its power spectral density (PSD; computed using fast Fourier transform) were manually selected to remove eye (1–15 Hz), muscle (110–140 Hz), and other noise artifacts^41^. A second round of automatic channel exclusion was performed to remove any channels exceeding two standard deviations of the new mean across channels. A second round of automatic trial exclusion was performed to remove any trials containing data exceeding five standard deviations of the moving average. Data were then manually inspected to remove any channels and trials with residual noise. Clean data were z-scored in the time domain to normalize amplitudes across sleep and wake states. Preprocessing was completed using functions from the FieldTrip toolbox for MATLAB^42^.

### High-frequency broadband activity

All analyses followed standards of HFB analysis in iEEG data^43^. HFB power was calculated from 70–150 Hz in steps of 10 Hz (i.e., 70–80, 80–90, … 140–150 Hz) using a multitaper frequency transformation based on discrete prolate spheroidal sequences (DPSS)^44^. The trial length was zero-padded up to the next power of 2 to minimize filter-induced artifacts. The data within each frequency band was normalized by frequency multiplication, and the average across frequencies was calculated in each channel.

### Statistical analyses

We ran linear mixed-effects models per channel, with the fixed effect of wake/sleep state and random effects of subject and trial nested in subject modeled as intercepts^45^, FDR-corrected for multiple comparisons across channels^46^. Linear mixed-effects models better account for nuisance variables than ANOVA and linear regression, and they allow for the inclusion of both subjects and states with few and/or unequal numbers of trials^47^. We first ran the models using all trials and subjects.

To determine how the number of trials influenced the reliability of results, we repeated these analyses while varying the number of trials included per subject (1–20 trials). We repeated analyses 10 times per trial count, randomly selecting a different set of trials for each replication. We defined reliability for each channel as the percentage of analyses within a given trial count that revealed greater HFB power during wake versus sleep states.

To determine how the number of subjects influenced reliability, we repeated the analyses with 1–20 trials per subject and additionally varied the number of subjects included (1–18 subjects). We randomly selected a different set of subjects for each subject count and used the same subjects while varying the number of trials used for each subject. We repeated analyses 10 times per trial and subject count, randomly selecting a different set of trials for each replication. We defined reliability for each channel as the percentage of analyses within a given trial and subject count that revealed greater HFB power during wake versus sleep states.

To further explore the reliability of the group-level results, we used the READIE Toolbox to calculate internal consistency reliability, which captures how correlated trials are to one another^27^. To do this, we calculated the difference in HFB power between states (wake - sleep) for pairs of trials in each subject, in temporal order. This yielded one difference value per trial pair, to be treated as a trial. Unpaired trials were excluded from analysis. From this list of HFB power differences, we calculated internal consistency reliability for each channel with split-half reliability. For each of 1000 iterations, we randomly subsampled 2–20 trials (i.e., difference scores) in steps of 2. Thresholds for internal consistency reliability were defined by the READIE Toolbox as acceptable (.60), good (.70–.80), and excellent (.90).

To determine whether the significant group-level wake > sleep effects can be detected at an individual level, we ran linear mixed-effects models separately for each subject, with the fixed effect of wake/sleep state and random effect of trial. As the wake > sleep effect seen in intracranial data is not hemisphere specific^8^, we ran these models on the mean HFB of the channels located on the midline or bilaterally that had a wake > sleep effect in the group analyses (Fz, Cz, Pz, C3, C4, O1, O2). Similarly to the group data, to determine how the number of trials influenced the reliability of results, we repeated these analyses while varying the number of trials included per subject (1–20 trials). We repeated the analyses 10 times per trial count, randomly selecting a different set of trials for each replication. We defined reliability for the mean HFB power across channels as the percentage of analyses within a given trial count that revealed greater HFB power during wake versus sleep states.

## Acknowledgements

We thank Kristopher Anderson, Kurtis Auguste, and Maya Cano for their help in data acquisition and initial preprocessing. This research was funded by grants from the National Institute of Neurological Disorders and Stroke (T32NS047987 to A. M. H. and J. B. Y., R00NS115918 to E. L. J., R01NS021135 to R. T. K.), This research was supported in part through the computational resources and staff contributions provided for the Quest high performance computing facility at Northwestern University which is jointly supported by the Office of the Provost, the Office for Research, and Northwestern University Information Technology.

## References

1. Leonard, M.K. et al. Nature 626, 593–602 (2024).

2. Leszczyński, M. et al. Sci. Adv. 6, eabb0977

3. Logothetis, N.K., Pauls, J., Augath, M., Trinath, T. & Oeltermann, A. (2001).

4. Manning, J.R., Jacobs, J., Fried, I. & Kahana, M.J. J. Neurosci. 29, 13613–13620 (2009).

5. Ray, S., Crone, N.E., Niebur, E., Franaszczuk, P.J. & Hsiao, S.S. J. Neurosci. 28, 11526–11536 (2008).

6. Rich, E.L. & Wallis, J.D. Nat. Commun. 8, (2017).

7. Watson, B.O., Ding, M. & Buzsáki, G. Eur. J. Neurosci. 48, 2482–2497 (2018).

8. Gross, D.W. & Gotman, J. Neuroscience 94, 1005–1018 (1999).

9. Johnson, E.L., Kam, J.W.Y., Tzovara, A. & Knight, R.T. J. Neural Eng. 17, (2020).

10. Parvizi, J. & Kastner, S. Nat. Neurosci. 21, 474–483 (2018).

11. Gerner, N., Thomschewski, A., Marcu, A., Trinka, E. & Höller, Y. Front. Neurol. 11, 432 (2020).

12. Höller, P., Trinka, E. & Höller, Y. Comput. Intell. Neurosci. 2018, 1–9 (2018).

13. Subramanian, A.K. et al. bioRxiv 2025.04.07.647612 (2025).doi:10.1101/2025.04.07.647612

14. Ball, T. et al. NeuroImage 41, 302–310 (2008).

15. Onton, J. & Makeig, S. Front. Hum. Neurosci. 3, 1–18 (2009).

16. Zelmann, R., Lina, J.M., Schulze-Bonhage, A., Gotman, J. & Jacobs, J. Brain Topogr. 27, 683–704 (2014).

17. Johnson, E.L., Tang, L., Yin, Q., Asano, E. & Ofen, N. Sci. Adv. 4, eaat3702 (2018).

18. Johnson, E.L. & Knight, R.T. Intracranial EEG 143–154 (2023).doi:10.1007/978-3-031-20910-9_10

19. Ofen, N., Tang, L., Yu, Q. & Johnson, E.L. Dev. Cogn. Neurosci. 36, 100613 (2019).

20. Yin, Q., Johnson, E.L. & Ofen, N. Dev. Cogn. Neurosci. 64, 101312 (2023).

21. Ho, A.L. et al. Neurosurg. Focus 45, E7 (2018).

22. Kiesler, J. & Ricer, R. Am. Fam. Physician 67, 2547–2552 (2003).

23. Duc, G. & Largo, R.H. Pediatrics 78, 904–908 (1986).

24. D’Antoni, A.V. et al. Childs Nerv. Syst. 33, 909–914 (2017).

25. Katz, J., Armstrong, C., Kvint, S. & Kennedy, B.C. Epilepsy Behav. Rep. 19, 100552 (2022).

26. Voytek, B. et al. J. Cogn. Neurosci. 22, 2491–2502 (2010).

27. Xu, W. et al. Dev. Cogn. Neurosci. 70, 101458 (2024).

28. Lipsett, B.J., Reddy, V. & Steanson, K. StatPearls (2025).

29. Kabdebon, C. et al. NeuroImage 99, 342–356 (2014).

30. Skelton, A.E., Maule, J. & Franklin, A. Child Dev. Perspect. 16, 90–95 (2022).

31. Alho, K., Sainio, K., Sajaniemi, N., Reinikainen, K. & Näätänen, R. Electroencephalogr. Clin. Neurophysiol. Potentials Sect. 77, 151–155 (1990).

32. Bishop, D.V.M., Hardiman, M.J. & Barry, J.G. Dev. Sci. 14, 402–416 (2011).

33. Edwards, E., Soltani, M., Deouell, L.Y., Berger, M.S. & Knight, R.T. J. Neurophysiol. 94, 4269–4280 (2005).

34. Clayson, P.E. & Miller, G.A. Int. J. Psychophysiol. 111, 68–79 (2017).

35. Meyer, A., Bress, J.N. & Proudfit, G.H. Psychophysiology 51, 602–610 (2014).

36. Mon, S.K., Manning, B.L., Wakschlag, L.S. & Norton, E.S. Dev. Cogn. Neurosci. 70, 101459 (2024).

37. Odabaee, M. et al. NeuroImage 96, 73–80 (2014).

38. Faix, R.G. J. Pediatr. 100, 304–306 (1982).

39. Electroencephalogr. Clin. Neurophysiol. 10, 370–375 (1958).

40. Klem, G.H., Lüders, H.O., Jasper, H.H. & Elger, C. Electroencephalogr. Clin. Neurophysiol. Suppl. 52, 3–6 (1999).

41. Hipp, J.F. & Siegel, M. Front. Hum. Neurosci. 7, (2013).

42. Oostenveld, R., Fries, P., Maris, E. & Schoffelen, J.M. Comput. Intell. Neurosci. 2011, (2011).

43. Mercier, M.R. et al. NeuroImage 260, 119438 (2022).

44. Mitra, P.P. & Pesaran, B. Biophys. J. 76, 691–708 (1999).

45. Yu, Z. et al. Neuron 110, 21–35 (2022).

46. Benjamini, Y. & Hochberg, Y. J. R. Stat. Soc. Ser. B Stat. Methodol. 57, 289–300 (1995).

47. Heise, M.J., Mon, S.K. & Bowman, L.C. Dev. Cogn. Neurosci. 54, 101070 (2022).

